# DHX15 inhibits mouse APOBEC3 deamination

**DOI:** 10.1101/2024.08.19.608612

**Authors:** Wenming Zhao, Susan R. Ross

**Affiliations:** Department of Microbiology and Immunology, University of Illinois Chicago College of Medicine, Chicago, IL 60612

## Abstract

APOBEC3 family proteins are critical host factors that counteract and prevent the replication of retroviruses and other viruses through cytidine deamination. Human APOBEC3 proteins inactivate HIV-1 through the introduction of lethal mutations to viral genomes. In contrast, mouse APOBEC3 does not induce DNA hypermutation of murine retroviruses, although it retains functional cytidine deaminase activity. Why mouse APOBEC3 does not effectively deaminate murine retroviruses is still unknown. In this study, we found that the dead box helicase DHX15 interacts with mouse APOBEC3 and inhibits its deamination activity. DHX15 was packaged into murine leukemia virus (MLV) virions independent of its binding with APOBEC3. Moreover, DHX15 knockdown inhibited MLV replication and resulted in more G-to-A mutations in proviral DNA. Finally, DHX15 knockdown induced DNA damage in murine cells, suggesting that it plays a role in preserving genome integrity in cells expressing mouse APOBEC3 protein.

## Introduction

Apolipoprotein B mRNA editing enzyme catalytic polypeptide 3 (APOBEC3) cytidine deaminases play important roles in the intrinsic response to virus infection (Stavrou & Ross, 2015). *APOBEC3* genes are highly diversified. The human genome encodes seven *APOBEC3* genes, while the mouse encodes a single gene (Salas-Briceno *et al*, 2020). Human APOBEC3G (hAPOBEC3G) was discovered because it restricts Viral Infectivity Factor (vif)-deficient HIV-1 (Sheehy *et al*, 2002). In cells infected with Vif-deficient HIV-1, APOBEC3G is packaged into progeny virions and acts in target cells, where it deaminates dC residues in virus minus strand DNA during reverse transcription, thus causing G to A hypermutation in newly synthesized plus strands (Kao *et al*, 2003; Marin *et al*, 2003; Sheehy *et al*, 2003a; Zhang *et al*, 2003). To counteract this, HIV Vif binds APOBEC3 in virus-producer cells and redirects it to degradation in the proteasome, preventing virion incorporation and protecting the viral genome from mutation (Mehle *et al*, 2004; Sheehy *et al*, 2003b; Yu *et al*, 2003). Several APOBEC3 proteins have also been implicated in genome mutation in cancer cells (Burns *et al*, 2013; Carpenter *et al*, 2023).

APOBEC3 proteins also inhibit HIV-1 replication by cytidine deaminase-independent means, such as blocking reverse transcription and integration (Bishop *et al*, 2008; Mbisa *et al*, 2010; Newman *et al*, 2005). Mouse APOBEC3 (mAPOBEC3) does not efficiently induce hypermutation of MLV or mouse mammary tumor virus (MMTV) reverse transcripts and inhibits virus replication by blocking reverse transcription (MacMillan *et al*, 2013; Nair *et al*, 2014; Rulli *et al*, 2008; Stavrou *et al*, 2014; Stavrou *et al*, 2018). However, mAPOBEC3 retains its deaminase activity on other substrates (MacMillan *et al*., 2013; Nair *et al*., 2014).

Murine retroviruses have several means of inhibiting mAPOBEC3. MLV encodes several anti-APOBEC3 proteins, including a glycosylated form of the Gag polyprotein termed glycoGag, which blocks mAPOBEC3’s access to the reverse transcription complex (Stavrou *et al*, 2013), while the MLV P50 viral protein, encoded by an alternatively spliced *gag* RNA, blocks virion packaging of mAPOBEC3 (Zhao *et al*, 2020). The MLV protease, which cleaves the Gag polyprotein, may cleave and inactivate mAPOBEC3 (Abudu *et al*, 2006). Finally, the rapid processivity of MMTV reverse transcriptase prevents APOBEC3 from accessing single-stranded reverse transcripts (Hagen *et al*, 2019). While these viral proteins counteract APOBEC3, none specifically inhibits its deaminase activity.

Another possibility for the lack of retroviral DNA deamination by mAPOBEC3 is that a host factor(s) inhibits deamination. To test this, we used immunoprecipitation (IP) of cells over-expressing mAPOBEC3 and hAPOBEC3G coupled with mass spectrometry (MS) to identify host interacting proteins. One of the candidates that bound both proteins was DEAH-box helicase 15 (DHX15). DHX15 is a ubiquitously expressed, highly conserved protein which functions in multiple biological processes, including RNA splicing and editing, and ribosome assembly and biogenesis (Chen *et al*, 2014; Memet *et al*, 2017; Studer *et al*, 2020; Yoshimoto *et al*, 2009). DHX15 also serves as a virus sensor through binding double-stranded RNA (dsRNA) from RNA viruses (Lu *et al*, 2014). Studies also suggest that DHX15 contributes to carcinogenesis in some cancer types or acts as a tumor suppressor gene in others (Fan *et al*, 2023; Ito *et al*, 2017; Xiao *et al*, 2016; Xie *et al*, 2019; Zhang *et al*, 2022a). Knockdown DHX15 in leukemia cells causes DNA damage and cell cycle arrest (Li *et al*, 2024; Wang *et al*, 2022).

Here we show that DHX15 binds mAPOBEC3 and inhibits its deaminase activity, thereby preventing it from mutating the MLV genome. We also found that mouse cells depleted for DHX15 show increased APOBEC3-dependent DNA damage, suggesting that DHX15 may protect the genome.

## Results and Discussion

### DHX15 interacts with mAPOBEC3 and hAPOBEC3G

To determine if there were host factors inhibiting mAPOBEC3, we did IP/MS with extracts from 293T cells transfected with mAPOBEC3 and hAPOBEC3G FLAG-tagged expression vectors; a FLAG-tagged Stimulator of Interferon Genes (STING) expression vector served as a control. Coomassie blue staining revealed three bands in both the mAPOBEC3- and hAPOBEC3G- but not STING-expressing or untransfected cell lysates (bands 2, 3 and 4; Fig 1A). These bands contained 3 ribonuclear proteins: hnRNPK (2), hnRNPA1 (4) and Y-box binding protein 1 (YB1) (3). YB1 and hnRNP proteins were previously shown to bind APOBEC3G, validating our approach (Gallois-Montbrun *et al*, 2008; Gallois-Montbrun *et al*, 2007; Kozak *et al*, 2006). We also found one interacting band in the mAPOBEC3- but not the hAPOBEC3G-expressing samples. MS analysis showed that this band was DHX15.

**Fig 1.**
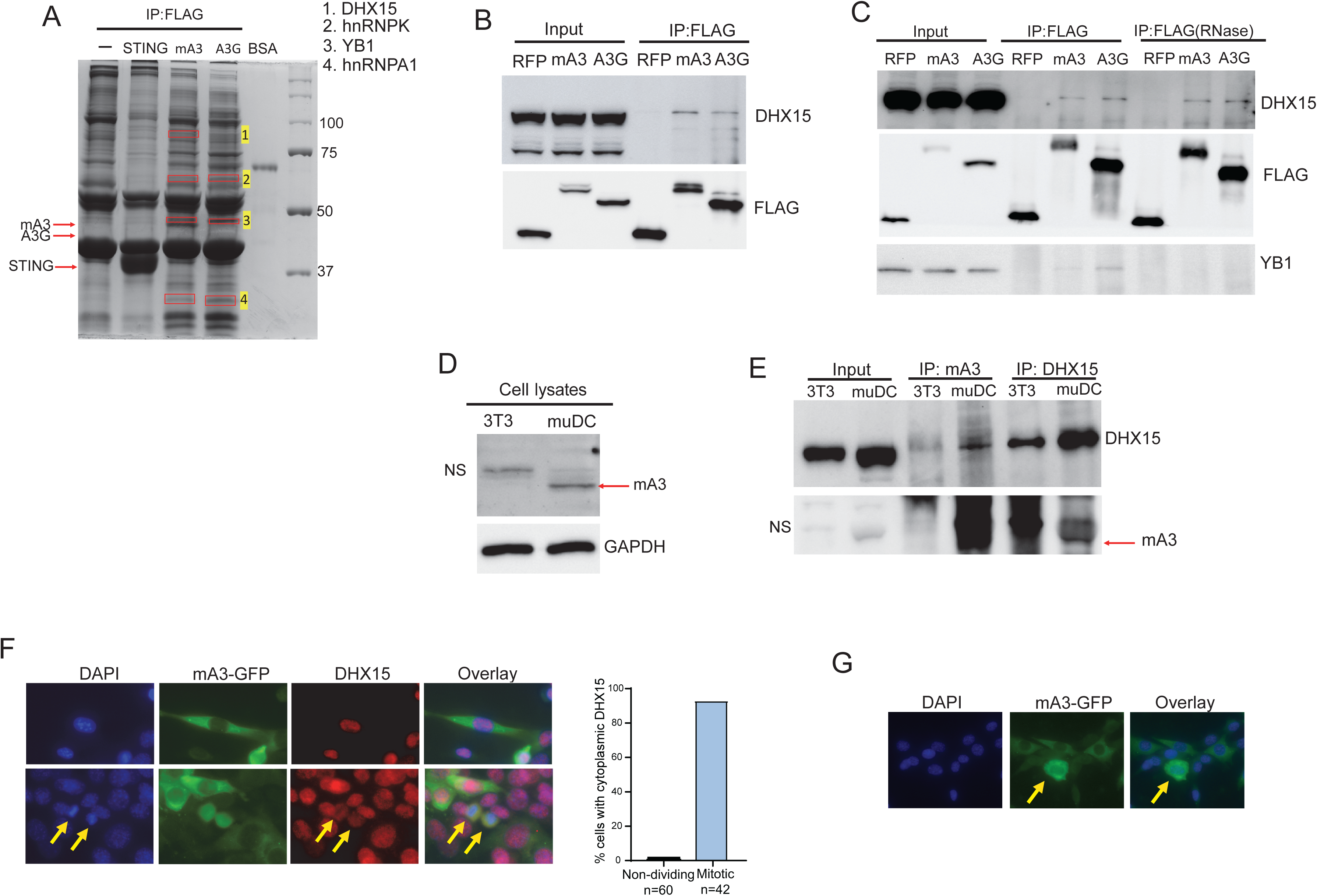
APOBEC3 and DHX15 interact. A) Coomassie-blue stained gels of co-immunoprecipitated extracts from 293T cells transduced with STING, mAPOBEC3 or hAPOBEC3G expression plasmids. Numbers correspond to the major proteins identified in the bands by MS. B) Co-IP of APOBEC3 proteins and DHX15 in 293T cells co-transfected with FLAG-tagged APOBEC3 or RFP expressing plasmids. C) Co-IP of APOBEC3 proteins and DHX15 or YB1. Prior to the IP, the lysates were treated with RNase A. D) Western blots of NIH3T3 and muDC cells probed for mAPOBEC3. NS, nonspecific band. E) Co-IP of mAPOBEC3 protein and DHX15. Lysates were immune precipitated, and blots were probed with anti-DHX15 and -mA3 antibodies. Shown is a representative Western blot of three different experiments. F) Colocalization of DHX15 and mouse A3 in dividing cells. NIH3T3 cells stably expressing GFP-tagged mouse APOBEC3 were fixed and stained with anti-DHX15 antibody (red) and DAP. The two yellow arrows indicated newly dividing cells. Shown to the right is quantification cells positive for cytoplasmic DHX15. G) Subcellular localization of GFP-mA3 in the stable cell line. Yellow arrow indicates nuclear staining.

To verify DHX15 binding to APOBEC3, we performed co-IP and Western blot assays with extracts from cells over-expressing mAPOBEC3, hAPOBEC3G or RFP as a control. Endogenous DHX15 bound both mAPOBEC3 and APOBEC3G but not RFP (Fig. 1B). However, APOBEC3G was more highly expressed than mAPOBEC3 yet pulled down less DHX15, suggesting that the interaction might be weaker (Fig. 1B and 1C). This may explain why DHX15 was not identified in the APOBEC3G IP/MS (Fig. 1A).

APOBEC3 and DHX15 are both RNA binding proteins. It was possible that the two proteins co-immunoprecipitated because of their RNA binding activity. We RNase A-treated the extracts prior to IP and found that mAPOBEC3 or hAPOBEC3G and DHX15 interaction did not depend on RNA, since RNase A treatment had no effect (Fig. 1C). In contrast, co-IP of mAPOBEC3 or hAPOBEC3G with YB-1 was ablated by RNaseA, as previously shown (Gallois-Montbrun *et al*., 2007).

We then examined endogenous DHX15 and mAPOBEC3 interaction using NIH3T3 cells, which don’t express mAPOBEC3, and the murine dendritic cell line MutuDC1940 (muDC), which expresses mAPOBEC3 (Fig. 1D). When anti-mAPOBEC3 antibody was used for the IPs, DHX15 could be detected in muDC but not NIH3T3 extracts (Fig. 1E). The reciprocal IP (anti-DHX15 IP, WB with anti-mAPOBEC3) also demonstrated that the two proteins interact (Fig. 1E).

DHX15 is found primarily in the nucleus, although it localizes to the cytoplasm during virus infection (Mosallanejad *et al*, 2014; Pattabhi *et al*, 2019; Zhang *et al*, 2022b). In contrast, mAPOBEC3 is largely cytoplasmic (Zhao *et al*., 2020). To determine if mAPOBEC3 and DHX15 co-localized in cells, we generated NIH3T3 cells stably expressing a green fluorescent protein (GFP)- tagged mAPOBEC3 and immunostained the cells with anti-DHX15 antibodies. As previously reported, mAPOBEC3 was mainly located in the cytoplasm, although in highly expressing cells it was also found in the nucleus (Fig. 1F, 1G). DHX15 was mostly nuclear, but relocalized from the nucleus to the cytoplasm in mitotic cells where it co-localized with mAPOBEC3 (Fig. 1F).

### DHX15 is packaged into virions

APOBEC3G is packaged into retrovirions through interaction with viral RNA and nucleocapsid (Alce & Popik, 2004; Khan *et al*, 2005; Okeoma *et al*, 2007; Schafer *et al*, 2004; Zennou *et al*, 2004). mAPOBEC3 is also incorporated into MLV virions (Stavrou *et al*., 2013; Zhang *et al*, 2008). We next tested whether DHX15 was packaged into MLV virions and whether packaging depended on mAPOBEC3 interaction. Wild type (APO+/+) and knockout (APO-/-) pups were infected with Moloney MLV and at 16 dpi, virions isolated from spleens were analyzed by western blot. Similar amounts of DHX15 protein were packaged into virions whether mAPOBEC3 was present or not (Fig. 2A). Thus, DHX15 protein is packaged into MLV virions independent of mAPOBEC3 binding. The block to deamination of reverse transcripts in target cells may occur when DHX15 re-localizes to the cytoplasm during cell division, or because virus infection causes its relocation, as previously described (Mosallanejad *et al*., 2014; Pattabhi *et al*., 2019; Zhang *et al*., 2022b).

**Fig 2.**
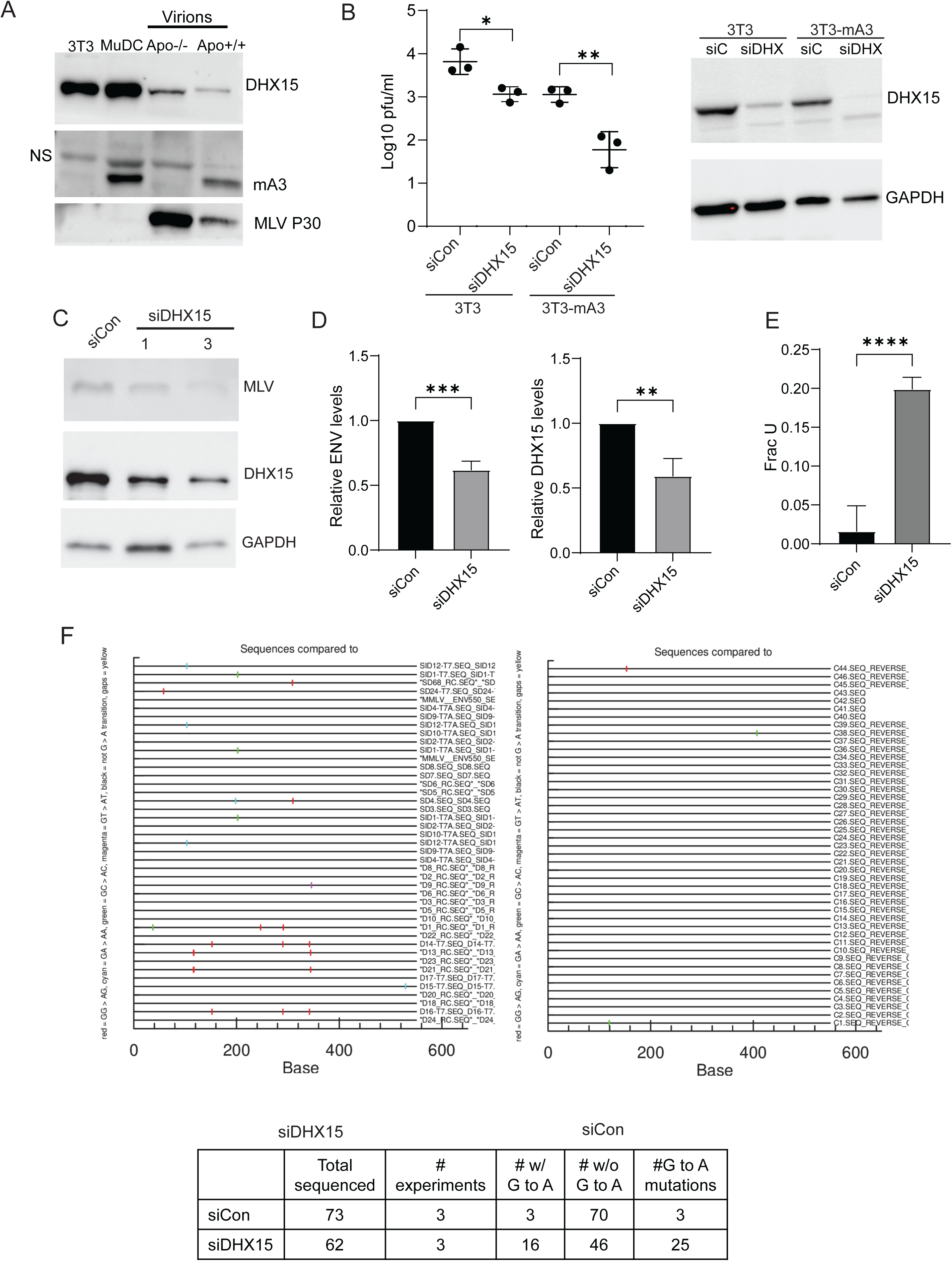
DHX15 inhibits mAPOBEC3 deamination of MLV. A) Western blot of MLV virions isolated from APO-/- and APO+/+ mice infected with MLV virus and probed with anti-DHX15 (upper) or anti-MLV (lower) antisera. Shown is a representative blot of three different experiments. B) Virus titers (left panel) of the supernatants from infected cell lines with DHX15 siRNA or control (siCon) knockdown. Bar shows the average ± SD of 3 independent experiments, represented by individual points. *, *P* < 0.05; **, *P* < 0.01. Western blot (right panel) of the cell lysates from this experiment probed with anti-DHX15. Shown is a representative blot. C) Western blot of muDC cell lysates after DHX15 siRNA knockdown. Shown is a representative blot. D) Left panel: RT-qPCR with MLV SU-MLV primers levels in MLV-infected muDC cells line infected after DHX15 siRNA knockdown. Right panel: DHX15 knockdown levels. Shown is the average ± SD of 3 independent experiments. **, *P* < 0.01; ***, *P* < 0.001. E) Ex-qPCR method with SU-MLV primers was used to determine the fraction of uracils in MLV proviruses in infected muDC cells. Shown is the average ± SD of 4 independent experiments. ****, *P* < 0.0001. F) Genomic DNA was isolated from the infected muDC cells described in D) was cloned and sequenced. Shown is the analysis of 3 independent experiments. Shown are the G to A changes in the sequences (red bars).

### DHX15 knockdown inhibits MLV infection and increases G to A mutation in proviral DNA

We then examined whether DHX15 affected mAPOBEC3’s ability to inhibit MLV infection. NIH3T3 cells stably expressing either GFP-tagged mAPOBEC3 or GFP alone were transfected with DHX15 siRNA (siDHX15) or scrambled control siRNA (siCon) and infected with MLV. Virus titers were determined at 72 hrs post-infection. DHX15 knockdown slightly reduced virus titers in NIH3T3 cells (Fig. 2B). However, in mAPOBEC3-expressing cells, DHX15 reduced virus titers more than 10-fold (Fig. 2B).

mAPOBEC3 primarily restricts MLV by cytidine-deaminase-independent means, although it retains functional enzymatic activity (Nair *et al*., 2014; Rulli *et al*., 2008; Stavrou *et al*., 2018). To determine whether DHX15 protein affected mAPOBEC3-mediated deamination, MLV-infected muDC cells were treated with control or DHX15 siRNAs. At 48 hrs post-infection, protein and RNA analysis showed that MLV Env and DHX15 protein levels were reduced by DHX15 knockdown (Fig. 2C). Env RNA was also reduced by about 50%, similar to the reduction in level of viral protein (Fig. 2D). Genomic DNA from the infected DHX15-depleted muDC cells was used to determine uracil incorporation in proviral DNA. Uracil levels were dramatically higher in proviral DNA isolated from DHX15-depleted cells (Fig. 2E).

We next tested whether DHX15 in mAPOBEC3-expressing cells resulted in increased G-to-A mutations. An *env* gene segment from MLV-infected muDC cells was sequenced (Stavrou *et al*., 2014). DHX15-depletion resulted in increased numbers of G-to-A mutations in this fragment (Fig. 2F). The more dramatic effect on uracil incorporation compared to G-to-A mutations likely reflects uracil deglycosylase (UNG) activity in base excision repair; we showed previously that UNG repairs APOBEC3-mediated mutations in proviruses (Salas-Briceno & Ross, 2021).

### DHX15 interact withs mAPOBEC3’s N-terminus

mAPOBEC3 protein has two conserved zinc-coordinating cytidine deaminase (CD) domains (Harris & Dudley, 2015). The N-terminal CD1 encodes the deaminase, while the C-terminal CD2 is essential for encapsidation (Browne & Littman, 2008; Hakata & Landau, 2006). To determine which domain interacted with DHX15, we subcloned full-length and N-terminal and C-terminal domains into FLAG-tagged expression vectors; FLAG-tagged RFP served as a control (Fig. 3A). These were transiently transfected with the DHX15 vector into 293T cells and co-IPs were performed. While DHX15 strongly interacted with the N-terminal domain of mAPOBEC3, showing binding comparable to the full-length protein, the C-terminal domain interaction was weak (Fig. 3B).

**Fig 3.**
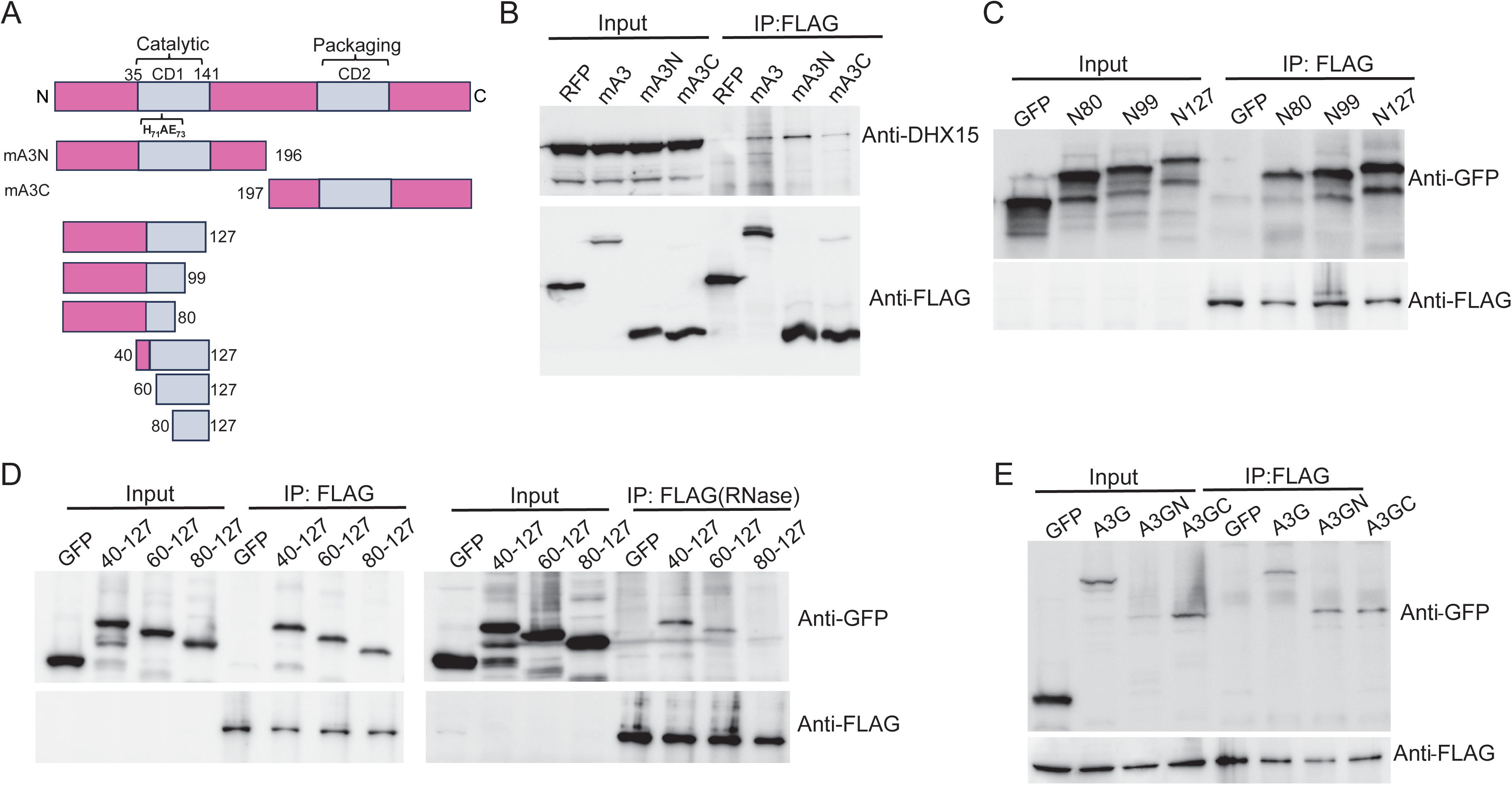
mAPOBEC3 and hAPOBEC3G CD domains interact with DHX15. A) Diagrams of expression constructs. B) Co-IP of different domains of mAPOBEC3 and DHX15. 293T cells were transfected with FLAG-tagged RFP, mA3, mA3N or mA3C expressing plasmids, the lysates were immunoprecipitated and western blots were probed with anti-DHX15 and –FLAG antibodies. C) and D) Co-IP of mAPOBEC3 CD1 truncations and DHX15. D) Co-immunoprecipitations were carried out without (right panel) and with RNaseA (left) treatment prior to immunoprecipitation. Shown is a representative Western blot of three different experiments. E) Co-IP of hAPOBEC3G N and C domains and DHX15. 293T cells were co-transfected with FLAG-tagged DHX15 and GFP tagged A3G, A3GN or A3GC expressing plasmids. Lysates were immunoprecipitated and western blots were probed with anti-FLAG and –GFP antibodies.

We next generated a series of mAPOBEC3 CD1 deletions (Fig. 3A). Co-IP experiments showed that fragments containing up to 80 amino acids of the N terminus bound DHX15 (Fig. 1C), as well as fragments containing N40-N127, N60-N127 and N80-N127 (Fig. 1D). To determine if DHX15 and the mAPOBEC3 truncations co-immunoprecipitated because both bound RNA, we also treated the cell lysates with RNaseA. RNaseA abolished the interaction between DHX15 and mAPOBEC3 N80-N127, but not between N40-N127 and N60-N127 and DHX15 (Fig. 3D). Therefore, the strongest interaction site is within the N60-N80 AA region, which contains the deaminase domain active site motif His_71_-X-Glu_73_ (Fig. 3A).

hAPOBEC3G also has two CD domains, although its deaminase activity is in the C-terminal CD2, while packaging and homodimerization are mediated by the N-terminal CD1 (Hakata & Landau, 2006). The hAPOBEC3G N- and C-terminal domains were GFP-tagged and co-transfected into 293T cells with FLAG-tagged DHX15. Similar to mAPOBEC3, the DHX15 interacted with the full-length, as well as the N- and C-terminal domains of APOBEC3G, although the N-terminal and C-terminal domains did not show a strong difference in binding (Fig. 3D).

### DHX15 inhibits mAPOBEC3 deamination

To investigate whether DHX15 inhibits mAPOBEC3 deamination, we depleted DHX15 by siRNA knockdown in mouse embryo fibroblasts (MEFs) isolated from wild-type (APO+/+) and knockout (APO-/-) mice. Following knockdown, we measured deamination activity using the Epigenase APOBEC3 Cytidine Deaminase Activity Assay Kit. DHX15 knockdown led to a significant increase in deamination activity in APO+/+ MEFs, but no significant change in deamination activity in APO-/- MEFs, demonstrating DHX15’s inhibitory effect on mAPOBEC3 (Fig 4A).

**Fig 4.**
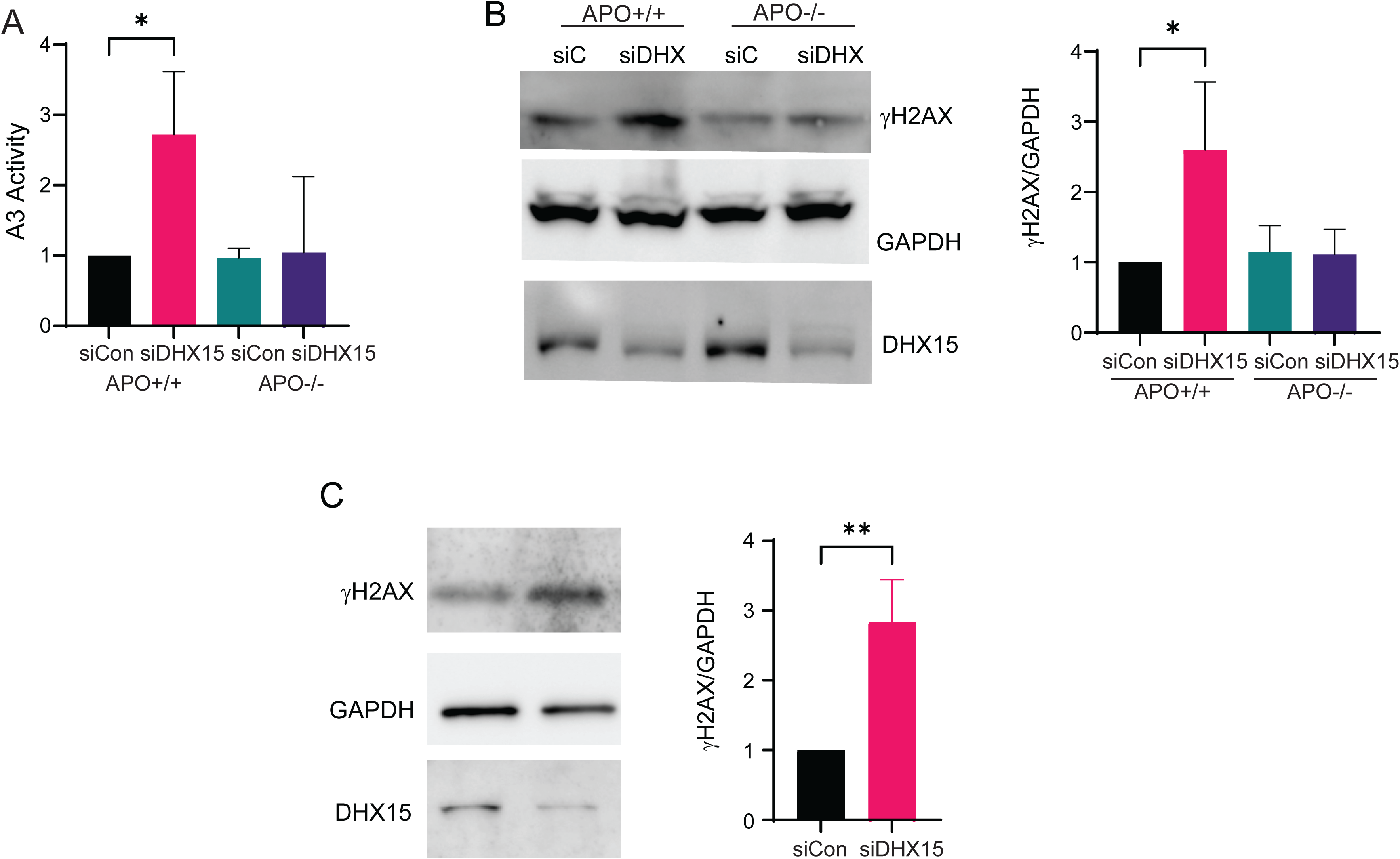
DHX15 knockdown induces DNA damage. A) In vitro deamination activity assay of MEFs isolated from both APOBEC3 wild-type (APO+/+) and knockout (APO-/-) mice after DHX15 knockdown. Shown is the average ± SD of 3 independent experiments. *, *P* < 0.05. B) Left panel: western blot of MEF cell lysates after DHX15 siRNA knockdown. Anti-p-γH2AX, anti-GAPDH and anti-DHX15 antibodies were used. Shown is a representative blot. Right panel: quantification of p-γH2AX protein levels. Shown is the average ± SD of 3 independent experiments. *, *P* < 0.05. C) Western blot (left panel) of the cell lysates of muDC cells after DHX15 knockdown. Anti -p-γH2AX, anti-GAPDH and anti-DHX15 antibodies were used. Shown is a representative blot. (Right panel) Quantification of p-γH2AX protein level in muDCs knockdown DHX15 by siRNA (siDHX15) or control (siCon). Shown is the average ± SD of 3 independent experiments. **, *P* < 0.01.

### DHX15 knockdown induces genomic DNA damage

Previous studies showed that DHX15 deficiency leads to DNA damage in human leukemia cells (Wang *et al*., 2022). We next performed DHX15 knockdown in APO+/+ and APO-/- MEFs mice and assessed DNA damage by measuring the levels of the DNA damage marker phosphorylated histone H2A.X (p-γH2AX). DHX15 knockdown significantly increased p-γH2AX levels in APO+/+ MEFs compared to APO-/- MEFs, indicating higher levels of mAPOBEC3-dependent DNA damage (Fig. 4B).

We performed a similar experiment with muDCs and found that DHX15 knockdown significantly elevated p-γH2AX levels compared to the control group (Fig. 4C). These results suggest that DHX15 plays a critical role in maintaining genomic integrity in mAPOBEC3-expressing murine cells.

Multiple experimental systems have shown that mAPOBEC3 does not efficiently deaminate murine retroviruses like MMTV and MLV, although *in vitro* assays show that it retains deamination activity (Bishop *et al*, 2004; MacMillan *et al*., 2013). Several mechanisms have been proposed for this inability to deaminate MLV, including exclusion of mAPOBEC3 from virions through the action of the viral p50 protein, blocking of mAPOBEC3 access to the reverse transcription complex by the glycoGag and mAPOBEC cleavage by MLV protease (Abudu *et al*., 2006; Stavrou *et al*., 2013; Zhao *et al*., 2020). Here, we show that a host protein, DHX15, inhibits mAPOBEC3’s ability to deaminate MLV reverse transcribed DNA.

DHX15 is an RNA helicase with multiple roles in RNA biology. It also acts as an RNA sensor and interacts with RIG-I, to limit infection by a number of RNA viruses, including SARS-CoV-2, picornaviruses, rhabodviruses and paramyxoviruses (Lu *et al*., 2014; Mosallanejad *et al*., 2014; Pattabhi *et al*., 2019; Xing *et al*, 2021; Zhang *et al*., 2022b). DHX15 has also been implicated in cancer, both through its role in splicing and because it binds proteins like c-MYC (Chen *et al*, 2018; Fan *et al*., 2023; Ito *et al*., 2017; Jing *et al*, 2018; Li *et al*., 2024; Xie *et al*., 2019). We now define an additional role for DHX15 as an inhibitor of the anti-retroviral cytidine deaminase APOBEC3.

DHX15 knockdown in mAPOBEC3-expressing cells decreased deaminase activity and induced more G-to-A mutations in MLV proviral DNA, suggesting that DHX15 inhibits mAPOBEC3 deamination. Interestingly, DHX15 bound mAPOBEC3’s CD1 domain, encoding the deaminase activity. These findings suggest that DHX15 plays a regulatory role in modulating mAPOBEC3 activity, potentially through direct interaction with cytidine deamination domain. It is possible that by incorporating DHX15 into virions, MLV utilizes this protein to protect itself from mAPOBEC3-mediated mutation. However, DHX15 does not apparently inhibit mAPOBEC3’s ability to block reverse transcription, which may be due to its weaker binding to the C terminal nucleic acid binding domain. Although there are multiple host mechanisms for restricting MLV, that the virus is able to partially overcome them likely reflects the long co-evolution of this virus and its host species.

Knockdown experiments targeting DHX15 demonstrated its role in inhibiting MLV replication. However, this inhibition coincided with an increase in DNA damage. Viruses like MLV induce type I interferons in mice and as mAPOBEC3 is an interferon-inducible gene, its protein levels increase during viral infection (Gao *et al*, 2013; Okeoma *et al*, 2009). Thus, DHX15 may prevent genomic DNA damage when APOBEC3 levels increase during infection. Our findings suggest that DHX15 may have evolved to limit the potential damage that APOBEC3 proteins can inflict on the genome when they are induced during virus infection.

### Materials and Methods Ethics statement

All mice were housed according to the policy of the Animal Care Committee of the University of Illinois at Chicago, and all studies were performed in accordance with the recommendations in the Guide for the Care and Use of Laboratory Animals of the National Institutes of Health. The experiments performed with mice in this study were approved by the committee (UIC ACC protocol #18-168).

### Cell culture and transfection

NIH3T3 and 293T cells were cultured in Dulbecco’s modified Eagle’s medium (DMEM) supplemented with 10% fetal bovine serum (FBS), L-glutamine, and penicillin/streptomycin. MuDC1940 cells were cultured in Iscove’s MDM supplemented with 8% FBS, L-glutamine, penicillin/streptomycin, 0.05 mM beta-mercaptoethanol, and 10 mM Hepes (Fuertes Marraco *et al*, 2012). For plasmid transfections, Lipofectamine 3000 and 2000 (Invitrogen) were used for NIH3T3 cells and 293T cells, respectively. For siRNA transfections, Lipofectamine RNAiMAX (Invitrogen) was used.

### MEF cultures

MEFs were obtained from day E17 to E18 APO+/+ and APO-/- fetuses. To generate MEFs, the heads and red organs of each embryo were removed, the carcasses were minced, and the lysates were incubated for 30min at 37 °C in 0.25% trypsin (Gibco). Trypsin was inactivated by adding complete DMEM, the lysates were centrifuged at 300 × g for 5 min, and the cell pellet was resuspended in complete DMEM and plated onto 10-cm cultures dishes.

### Mass spectrometry

HEK293T cells were transfected with plasmids expressing FLAG-tagged APOBEC3 proteins. At 24 h after transfection, cells were lysed in 1x cell lysis buffer (Cell Signaling Technology [CST], 9803) containing Halt protease and phosphatase inhibitor cocktail (Thermo Scientific). Supernatants were incubated with monoclonal anti-FLAG antibody (Sigma F3165) and then with protein A/G Plus-agarose (Santa Cruz Biotechnology). Proteins were eluted from the agarose and separated on 12% SDS polyacrylamide gels. After Coomassie blue staining and destaining, gel bands were treated with 50% acetonitrile, 25 mM ammonium bicarbonate (ABC), reduced with 10mM Dithiothreitol in 25mM ABC and alkylated with 50 mM IAA. After further washing with 25mM ABC dehydration in 100% acetonitrile, the pieces were re-hydrated in 50mM ABC containing 10 ug/mL Trypsin (Promega), followed by extraction in 50% acetonitrile/0.1 % formic acid (FA). This step was repeated two more times and the elutes were combined and dried. Upon reconstituting in 5% acetonitrile in 0.1% FA in water the samples were desalted, washed with 5% acetonitrile in 0.1% FA and eluted with 50% ACN, 0.1% FA. After drying, the peptides were suspended in 5% acetonitrile, 0.1% FA buffer for LC-MS. The digested peptides were analyzed using a Q Exactive HF mass spectrometer coupled with an UltiMate 3000 RSLC nanosystem with a Nanospray Frex Ion Source (Thermo Fisher Scientific). Digested peptides were loaded into a Waters nanoEase M/Z C18 (100Å, 5um, 180um x 20mm) trap column and then a 75 μm x 150mm Waters BEH C18(130A, 1.7um, 75um x 15cm) and separated at a flow rate of 300nL/min. Full MS scans were acquired in the Q-Exactive mass spectrometer over 374-1400 m/z range with a resolution of 120,000 (at 200 m/z) from 5 min to 45 min. The AGC target value was 3.00E+06 for the full scan. The 15 most intense peaks with charge states 2, 3, 4, 5 were fragmented in the HCD collision cell with a normalized collision energy of 28%; these peaks were then excluded for the 30s within a mass window of 1.2 m/z. A tandem mass spectrum was acquired in the mass analyzer with a resolution of 60,000. The AGC target value was 1.00E+05. The ion selection threshold was 1.00E+04 counts, and the maximum allowed ion injection time was 50 ms for full scans and 50 ms for fragment ion scans. Spectra were searched against the Uniprot human database using Mascot Daemon (2.6.0, updated on 08/11/20) with the following parameters; parent mass tolerance of 10 ppm, constant modification on cysteine alkylation, variable modification on methionine oxidation, deamidation of asparagine and glutamine. Search results were entered into Scaffold DDA software (v6.0.1, Proteome Software, Portland, OR) for compilation, normalization, and comparison of spectral counts.

### Co-immunoprecipitation and RNaseA treatment

Cells were lysed in cell lysis buffer (CST) containing Halt protease and phosphatase inhibitor cocktail (Thermo Scientific). Extracts were incubated with the indicated antibodies, followed by protein A/G Plus-agarose. The immunoprecipitated proteins were analyzed by immunoblotting using the indicated antibodies. For RNase A-treated samples, cell lysates were incubated with RNase A (50 µg/ml) at 37°C for 30 minutes prior to immunoprecipitation and then processed as described above.

### Western blot analysis

Affinity-purified polyclonal rabbit anti-mAPOBEC3 antibody has been previously described (Okeoma *et al*, 2010). Goat anti-MLV antibody (NCI Repository), rabbit anti-DHX15 (Invitrogen PA5-61413), rabbit anti-Phospho-Histone H2A.X (CST 9718S), anti-GAPDH (CST D16H11), mouse anti-FLAG (Sigma F3165) mouse anti-Myc (CST 2276), mouse anti-HA (CST 2367), rabbit anti-HA (Abcam 9110), rabbit anti-GFP (enQuire QAB10298), horseradish peroxidase (HRP)-conjugated anti-rabbit (CST 7074), anti-goat (Sigma-Aldrich A8919) and anti-mouse (Sigma-Aldrich A9044) antibodies were used for detection, using either Amersham ECL Prime Western blotting detection reagent (GE Healthcare Life Sciences) or Pierce ECL Western blotting substrate (Thermo Scientific).

### Virus isolation and virus titers

Moloney MLV was isolated from the supernatants of stably infected NIH3T3 cells (cells in which infection is allowed to spread to 100% of the culture and maintained in this state thereafter), as previously described (Stavrou *et al*., 2013). The medium was passed through a 0.45-μm filter and pelleted through a 25% sucrose cushion. MLV titers were determined by infectious center assay, as previously described (Low *et al*, 2009).

### *In vivo* infections

Tw-day-old mice were infected by intraperitoneal injection of 2x10^4^ ICs of MLV. Spleens were harvested at 16 dpi and virus was isolated, as previously described (Stavrou *et al*., 2013).

### Sequencing

DNA from MLV-infected muDC cells was isolated and a 549bp fragment from *env* was amplified using SU-MLV primers 5′-CCAATGGAGATCGGGAGACG-3′/5′-GTGGTCCAGGAAGTAACCCG-3′. The fragments were cloned into pCR2.1-TOPO vector (Invitrogen) and Sanger sequenced. Sequences were aligned using the MegAlign software, and G-to-A mutations were annotated by Hypermut (www.hiv.lanl.gov/contafent/sequence/HYPERMUT/hypermut.html).

### Deamination assay

The Epigenase™ APOBEC3 Cytidine Deaminase Activity/Inhibition Assay Kit was used for in vitro deamination assays, as recommended by the manufacturer (EpigenTek). Results were measured at 455 nm using an MCMI Biotek Synergy2 Plate Reader.

### Expression constructs

Expression plasmids pcDNA-FLAG and pcDNA-GFP were used were used to express FLAG tagged or GFP tagged A3 proteins, respectively. Cloning of pcDNA-FLAG-mA3, pcDNA-GFP-mA3, pcDNA- FLAG-mA3N and pcDNA-FLAG-mA3C were described previously (Zhao *et al*., 2020). The fragments encoding parts of mAPOBEC3 and hAPOBEC3G were amplified by PCR using the primers listed in Supplementary Table 1, then subcloned into pcDNA-GFP vector. All plasmids were sequenced prior to use.

### Stable cell line generation

NIH3T3 cells were transfected with pcDNA-GFP-mAPOBEC3 plasmids, then selected in DMEM medium containing 500 μg/ml G418(Sigma) for 2 weeks. The GFP-positive cells were sorted into single cells using a MoFlo Astrios cell sorter and cultured in DMEM containing 100 μg/ml G418 medium to generate stable cell lines. GFP-mAPOBEC3 expression was assessed by microscopy and Western blot analysis. Cells highly expressing GFP were selected for experiments.

### Immunofluorescence

Cells were seeded into 4-chamber culture slides (Millicell EZ slide; Millipore). The next day, cells were rinsed with ice-cold phosphate-buffered saline (PBS) and fixed with 4% paraformaldehyde for 15 min at room temperature, which was followed by permeabilization with 0.3% Triton X-100. The cells were subjected to immunofluorescence staining with anti-DHX15 antibody (Invitrogen PA6-61413) and Alexa 568-labeled anti-rabbit secondary antibody (Invitrogen, A-11011). The cells were examined by fluorescence microscopy (Keyence BZ-X710).

### Statistical analysis and data availability

Data shown are the averages of at least 3 independent experiments, or as otherwise indicated in the figure legends. Unpaired two-tailed t tests were performed using GraphPad Prism 10.1 software to calculate *P* values. All raw data have been deposited in the Mendeley data set found at: https://data.mendeley.com/datasets/kw6d5jvp6z/1.

### Conflict of Insurance

The authors declare that they have no conflict of interest.

## Acknowledgements

We thank David Ryan for help with the mice and the Mass Spectrometry Core in the Research Resources Center of the University of Illinois Chicago for MS analysis. The core is supported by NIH S10 shared instrumentation grant 1S10OD027016-01. Supported by NIH/NIAID R01 AI174538 and NIH/NIAID R01 AI085015.

---

**Supplementary Table 1.**
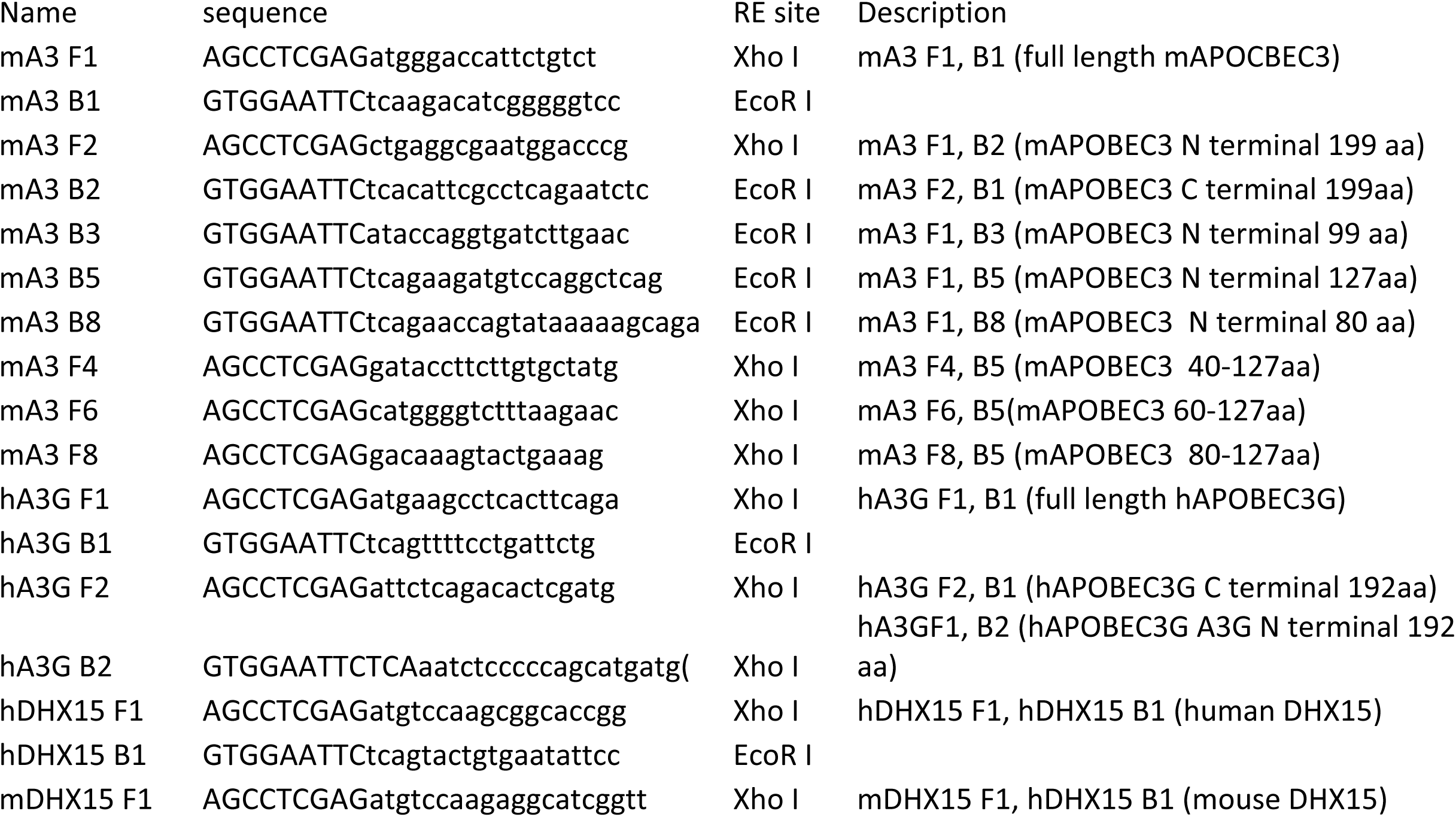
Primers used for cloning.

